# Label-free high-resolution infrared spectroscopy for spatiotemporal analysis of complex living systems

**DOI:** 10.1101/2023.01.05.522847

**Authors:** Nika Gvazava, Sabine Konings, Efrain Cepeda-Prado, Valeriia Skoryk, Chimezie H. Umeano, Jiao Dong, Iran A.N. Silva, Daniella Rylander Ottosson, Nicholas D. Leigh, Darcy E. Wagner, Oxana Klementieva

**Affiliations:** Department of Experimental Medical Science, Lund University; 22180 Lund, Sweden; MultiPark, Lund University; 22180 Lund, Sweden; NanoLund, Lund University; 22180 Lund, Sweden; Molecular Medicine and Gene Therapy, Department of Laboratory Medicine; 22184 Lund, Sweden; Lund Stem Cell Center, Lund University; 22100 Lund, Sweden; Wallenberg Centre for Molecular Medicine, Lund University; 22184 Lund, Sweden

## Abstract

Label-free chemical and structural imaging of complex living tissue and biological systems is the holy grail of biomedical research and clinical diagnostics. The current analysis techniques are time-consuming and/or require extensive sample preparation, often due to the presence of interfering molecules such as water, making them unsuitable for the analysis of such systems. Here, we demonstrate a proof-of-principle study using label-free optical photothermal mid-infrared microspectroscopy (O-PTIR) for fast, direct spatiotemporal chemical analysis of complex living biological systems at submicron resolution. While other analytical methods can provide only static snapshots of molecular structures, our O-PTIR approach enables time-resolved and in situ investigation of chemical and structural changes of diverse biomolecules in their native conditions. This comprises a technological breakthrough in infrared spectroscopy to analyze biomolecules under native conditions over time: in fresh unprocessed biopsies, living brain tissue, and vertebrates without compromising their viability.

**One-Sentence Summary:** Proof-of-principle application of non-destructive O-PTIR for high-resolution spatiotemporal chemical and structural analysis of unprocessed biopsies, living brain tissue, and vertebrates.

## Introduction

Spatiotemporal changes in the chemical and structural composition of cells underpin systemic health and disease. However, submicron-level chemical and structural alterations often occur before disease onset and before morphological changes can be detected using standard techniques at the tissue level such as through histology or immunohistochemical staining. Therefore, alternative techniques such as infrared (IR) imaging have been increasingly used in biomedical research and clinical diagnostics because they can simultaneously provide information about diverse biological macromolecules (e.g. proteins, lipids, metabolites, DNA, and RNA) as well as insights into their structural conformations (e.g. protein secondary structure^1^)^2^. Specifically, infrared microspectroscopy (μIR) has been used to characterize the compositional and molecular structural changes associated with the pathology of diverse diseases such as Alzheimer’s disease^3^, cancer^4^, septic arthritis^5^, and many others^6,7^. However, μIR analyses require extensive sample processing of biological materials due to the presence of water and the high degree of light scattering and absorbance which occurs in tissues. All biological tissues and organisms contain high levels of water which has strong IR absorption; thus, the detection of biological molecules is known to be extremely limited in aqueous environments. In addition, the high degree of light scattering in the majority of biological tissues limits the use of thick tissues in transmission-based IR approaches. Therefore, biological tissues must typically be dehydrated, chemically fixed and/or sectioned into thin slices for use with μIR based techniques which greatly adds both time and further variables to sample preparation and subsequently analysis. These are fundamental limitations which have thus far precluded the use of fresh, unprocessed biological tissue using μIR. Overcoming this limitation would open the door to unprecedented on-site rapid clinical diagnostics as well as novel applications in biomedical research.

## Results

Optical photothermal infrared spectroscopy (O-PTIR) is an emerging, novel IR-based technique that is currently revolutionizing spatial analysis of materials^8–17^. O-PTIR uses two co-linear beams of light (IR and a visible probe beam) to measure photothermal responses of the sample by assessing intensity changes of the reflected (or transmitted) probe beam at 532 nm. The fact that photothermal responses are measured with visible light wavelengths results in 5-10 times better resolution than traditional IR microspectroscopy techniques such as FTIR and QCL microscopies^8^. Importantly, when used in reflectance mode, it becomes a surface-scanning technique and therefore substrate chemistry (critical in transmission based IR methods) becomes negligible and perhaps more importantly, sample thickness becomes irrelevant^15^. While O-PTIR has been successfully used for the analysis of hydrated cells ^18^ and nematodes^8^, it is not yet known whether it can be applied for the analysis of living tissues and organisms without compromising their viability. One consideration is that O-PTIR employs high-intensity light beams that could impart thermal stress or light-induced injury, potentially altering the sample or impairing organism viability or development. However, this has not yet been tested. Here, we asked whether O-PTIR could be adapted for label-free rapid chemical and structural characterization of fresh, unprocessed tissue biopsies, including living brain tissue, as well as living vertebrates at a particularly vulnerable stage of their life cycle: during embryogenesis. We identified experimental imaging parameters that permit spectra acquisition and imaging of live tissues without compromising their viability by modifying the acquisition speed and laser power for use with the upright IR detector. We found that the combination of high-speed (1000 cm^-1^/s) laser scanning in combination with attenuated IR QCL / green light power (22 / 6 %) could be used to acquire high resolution spectra with the upright IR detector.

We first analyzed fresh mouse tissue biopsies from several organs (lung, heart, kidney, liver and salivary gland) and body parts (e.g. tail) by placing them on normal histological glass slides which were placed directly under the O-PTIR microscope (**Fig. 1A**). Despite using lower power laser intensities and high speed scan rates, the spectra obtained from hydrated tissues was of high quality and we observed the expected typical peaks in the Amide I /II regions corresponding to proteins, lipids, and phosphodiesters in all tissues analyzed (lung, heart, kidney, liver, salivary gland and skin)^19^ (shown by dashed lines in **Fig. 1B-F**). We noted distinct signal elevation at 1740 cm^-1^ in liver biopsies (**Fig. 1E**, orange trace), which corresponds to ester groups ((-C = O, -COOH) and is a well-known band used for conventional FTIR-based diagnosis of steatosis.^20,21^ The observed intensity changes may indicate different levels of lipidation^19^ or lipid oxidation^22^ and require complementary techniques for interpretation. Furthermore, we noticed elevations 1452 - 1396 cm^-1^ that can be assigned to in methylene groups of in proteins and lipids characteristic for salivary gland^23^. Our initial analysis demonstrated the utility of O-PTIR for direct label-free assessment of non-processed fresh biopsies immediately after excision and which demonstrated tissue specific IR bands previously identified to be unique in traditional IR approaches.

**Fig. 1.**
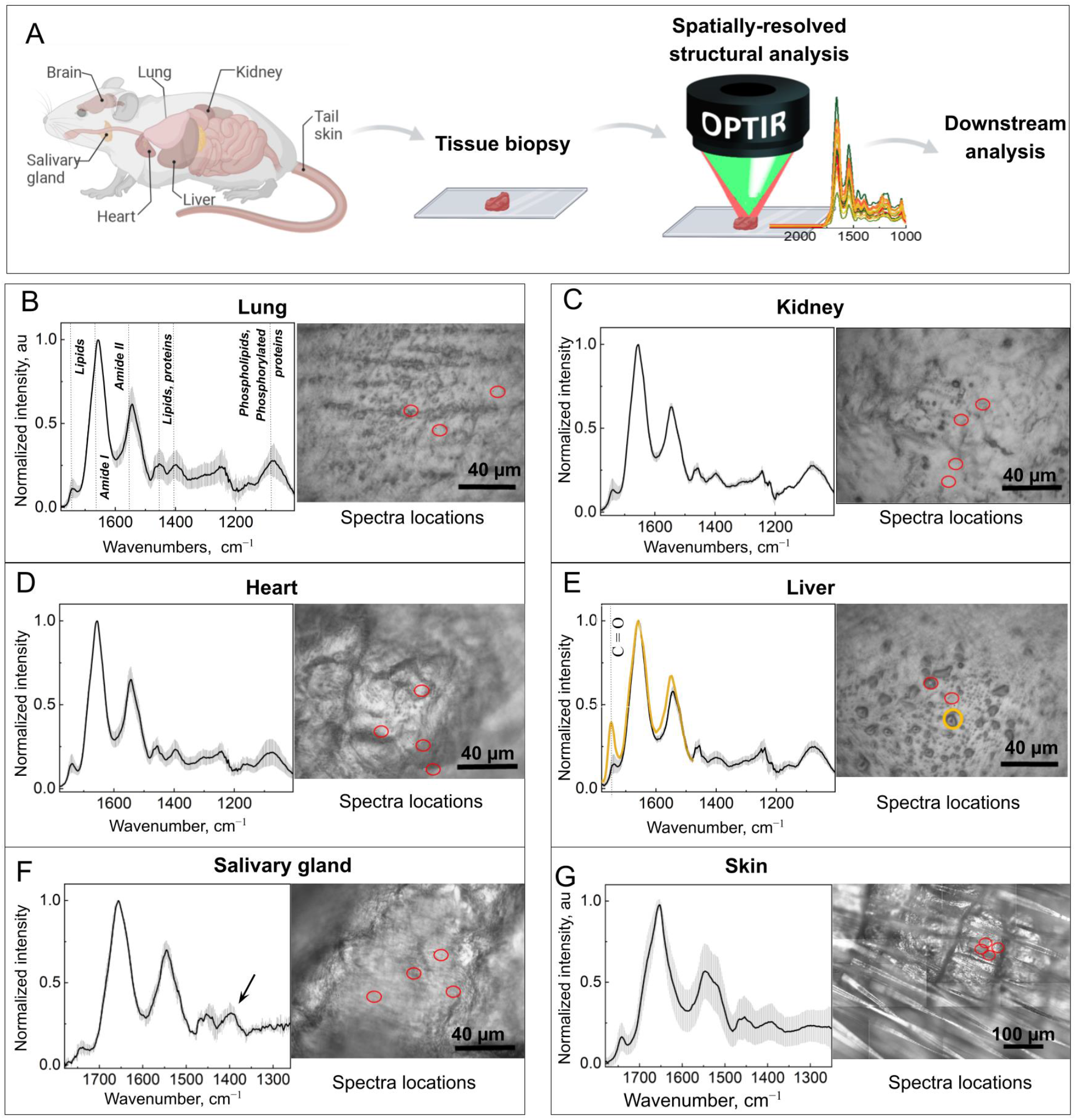
O-PTIR of fresh tissue biopsies. (**A**) Experimental flow. Created with BioRender.com. (**B–G**) Sample analysis. Averaged and normalized O-PTIR spectra and representative bright field images of the samples. Red circles indicate spectrum acquisition points, which were selected randomly for analysis. In (**E**), the yellow-colored spectrum trace was acquired at the location indicated by the orange circle and revealed a signal increase at 1740 cm^-1^, which suggests elevated ester levels. (**F, G**) Averaged and normalized O-PTIR spectra and representative bright field images of the indicated samples. The circles indicate spectrum acquisition points. The error bars represent s.d.; N=3–10 area from 3–4 tissue biopsies per organ.

While IR spectroscopy itself has not been reported to be destructive^1^, it is done on fixed and thus non-viable samples. On the other hand, O-PTIR requires high-intensity IR and visible light and one or both of these could potentially damage and/or affect sample viability. This is particularly important when subsequent analyses of the same sample are planned, e.g., for serial and time-resolved investigations. Therefore, we first assessed fresh biopsies which had been exposed to O-PTIR for evidence of structural alterations at the histological level. We found that all organs retained their metabolic activity after O-PTIR analysis (data not shown) and we did not observe any major tissue damage or signs of cellular necrosis after O-PTIR imaging (**Fig. S1)** further encouraging the use of O-PTIR either as a standalone technique or in combination with other complementary techniques.

As our findings using fresh tissue biopsies were highly encouraging as to the preservation of gross tissue morphology, we next optimized our setup to allow for imaging of brain slices and for serial imaging. In order to insure sufficient access to nutrients and to prevent dehydration over longer imaging times, we developed a sample support which allowed for brain slices of defined thickness to be imaged under aqueous conditions. Submerged tissue slices were placed into the sample support and covered by 0.35 mm CaF2 window transparent for both IR and 532 nm light; measurements were again performed using high-speed (1000 cm^-1^/s) and the upright IR detector with attenuated the IR QCL / green light power (22 and 6 %, respectively) to analyze spatiotemporal molecular changes in living brain tissue. We chose murine brain tissue due to the number of complementary tools and animal models which can be used to explore structural changes that are known to be important to human disease.

We first performed proof-of-principle experiments focusing on the detection of β-sheet structures in living brain tissue due to the fact that they have a specific IR signature between 1640 cm^-1^ and 1620 cm^-11,24^. To image β-sheet structures, we used brain slices generated from 12-month-old transgenic APP/PS1 mice, which express a chimeric mouse/human amyloid precursor protein and exhibit characteristic accumulations of β-sheet-folded amyloid-β proteins in the brain parenchyma, a known feature of Alzheimer’s disease ^25^. We first confirmed that O-PTIR imaging did not alter our ability to detect amyloid plaques in 12-month-old APP/PS1 brain slices. Following O-PTIR imaging, we stained APP/PS1 brain slices using Amytracker^R^ (EbbaBiotech, Solna, Sweden), a luminescent conjugated polyelectrolyte probe specific to amyloids^26^ and imaged using conventional and confocal fluorescence microscopy. We readily observed the presence of amyloid plaques following O-PTIR which we further confirmed by immunofluorescence staining using antibodies specific for amyloid plaques (**Fig. 2A-C, Supplemental Figure 2)**. Next, we evaluated whether O-PTIR affected metabolic activity or gross morphological tissue structure in APP/PS1 murine brains. We found that O-PTIR imaging of APP/PS1 brain slices did not affect metabolic activity using our O-PTIR imaging-based protocol or due to the *ex vivo* experimental conditions (**Fig. 2D**).

**Fig. 2.**
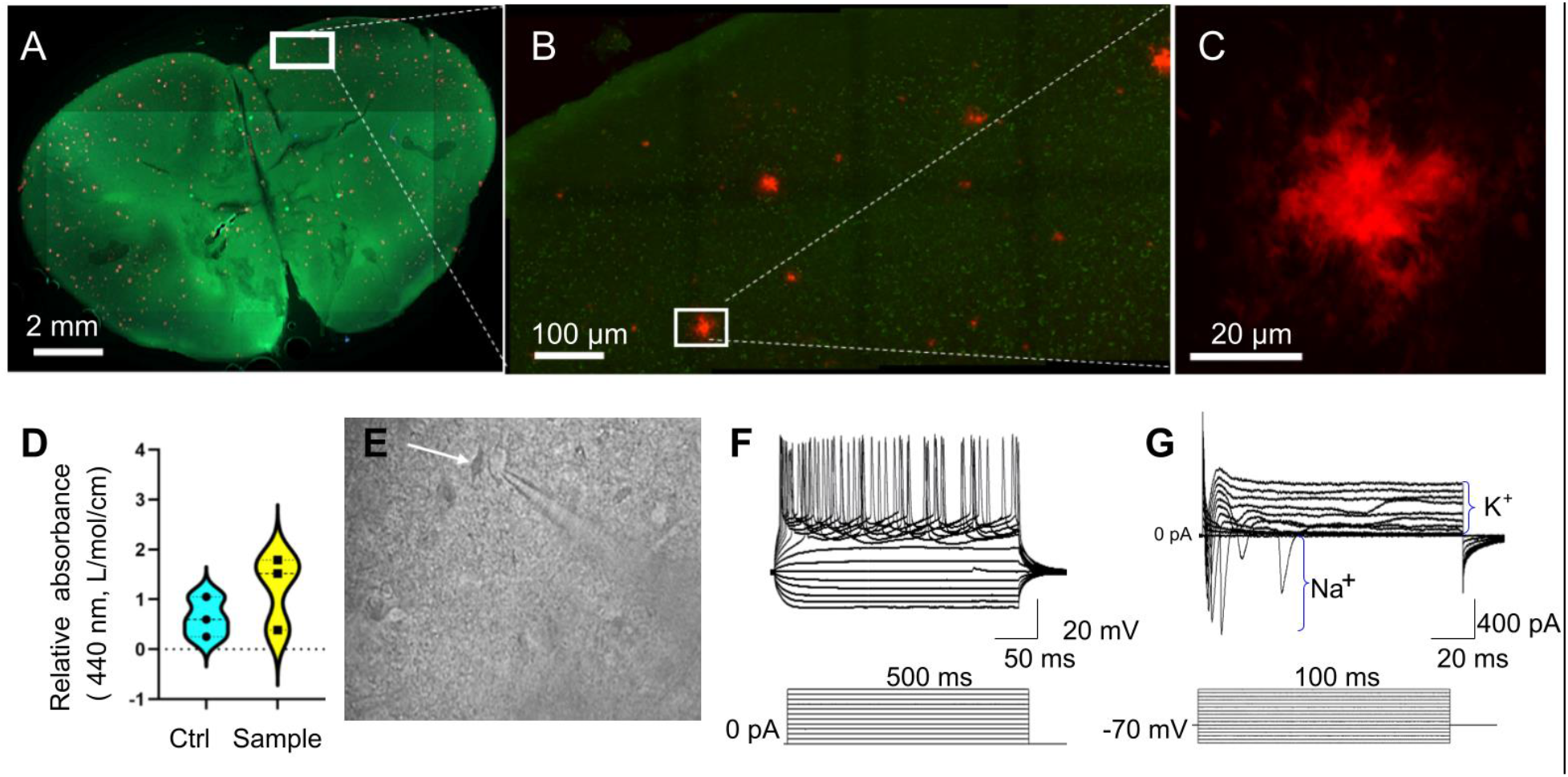
Brain tissue characterization for O-PTIR measurement. (**A**) Hydrated brain tissue was imaged after O-PTIR measurements using a Cytation5 multimode imager. Amyloid plaques were stained with Amytracker 520 after O-PTIR measurements and the tissue was fixed with 4 % PFA. White rectangle, O-PTIR measurement area. (**B**) Higher magnification image of the area in (**A**), with amyloid plaques stained in red. (**C**) High-resolution confocal imaging of amyloid plaque selected in B. (**D**) Brain tissue viability after O-PTIR measurement, as assessed using WST1. Control (Ctrl), the tissue that was incubated in the analysis buffer during O-PTIR analysis but that was not subjected to the analysis. (**E**) Bright field image of the brain tissue, measured with O-PTIR. The tissue was visualized with differential interface contrast imaging. The white arrow indicates a cortical neuron selected for whole-cell patch-clamp recording. (**F**) Induction of action potentials (firing pattern) to increasing current injection (from −100 pA to 200 pA with a delta of 20 pA for 500 ms, see below), recorded in current clamp mode at a holding potential of – 70 mV. (**G**) Activation of voltage-dependent sodium (Na+) and potassium (K+) channels using increasing depolarizing voltage steps from −70 mV to 40 mV with a delta of 10 mV for 100 ms in voltage clamp mode. Traces show inward and outward sodium and potassium currents over the membrane

As metabolic activity alone is not sufficient for assessing cellular level function after O-PTIR imaging, we next evaluated individual neuronal activity after O-PTIR imaging using electrophysiology with whole-cell patch-clamp electrophysiology recordings.^27^ We observed a robust firing pattern of action potentials as well as activation of voltage-gated sodium and potassium currents over the neuronal membrane, indicative of neuronal function after O-PTIR analysis at the single-cell level in the cortex, the location of O-PTIR imaging based measurements (**Fig. 2E-G, SI Fig. 2**). Collectively, these experiments confirmed that the neurons remain alive and functional within brain slices after 10-15 min of exposure to photothermal and high intensity visible light for IR spectroscopic analysis.

Having established a time window for O-PTIR measurements of living fully-hydrated brain tissue of APP/PS1 mice, we used O-PTIR to detect β-sheet structures in the native brain environment (**Fig. 3A-D**). We first located β-sheet amyloid plaques using fast hyperspectral array scanning over an extended area (10 spectra with 50 μm step, 1 sec per spectrum, over 1600 cm^-1^ 1700 cm^-1^ (**Fig. S2**), using the β-sheet associated IR absorption peak at 1630 cm^-1^ as an amyloid β-sheet marker^1,24^ and then acquired single spectra from individual plaques **(Fig. 3 E, F)**. Next, we took advantage of the superior submicron resolution capability of O-PTIR and our optimized protocol allowing for the use of O-PTIR in aqueous conditions to assess the distribution of β-sheet structures within individual amyloid plaques in hydrated tissue. We found that plaques exhibited little variability of β-sheet structures, as indicated by the O-PTIR acquired spectra **(Fig. 3 G)**. This result is in contrast to a large body of evidence suggesting that amyloid structures isolated from brain tissue are polymorphic^28–30^. However, all previous results have been collected exclusively with FTIR and from fixed and dehydrated tissue. Therefore, we next compared the O-PTIR acquired spectra of amyloid plaques in living brain tissue with parallel tissue slices from the same mouse that were chemically fixed with 4 % (v/v) paraformaldehyde (PFA) and dehydrated. Surprisingly, we observed an increase in the intensity of the β-sheet signal in fresh tissue as compared to the PFA-fixed tissue and a shift of the signal maximum from 1630 cm^-1^ in fresh tissue to 1634 cm^-1^ in PFA-fixed tissue (**Fig. 3 H, I**). In addition, we observed changes at the 1630 cm^-1^ bandwidth (**Fig. 3 J**), as detected from analysing the second derivative of IR spectra^31^. These striking observations of changes in β-sheet organization^32^ indicate that amyloid structures in living tissue can structurally differ from those that are chemically processed. This points to the opportunity and need to perform structural studies directly in living tissue, e.g., for the development of drugs that target β-sheet-enriched amyloids, such as lecanemab^33^ or aducanumab^34^.

**Fig. 3.**
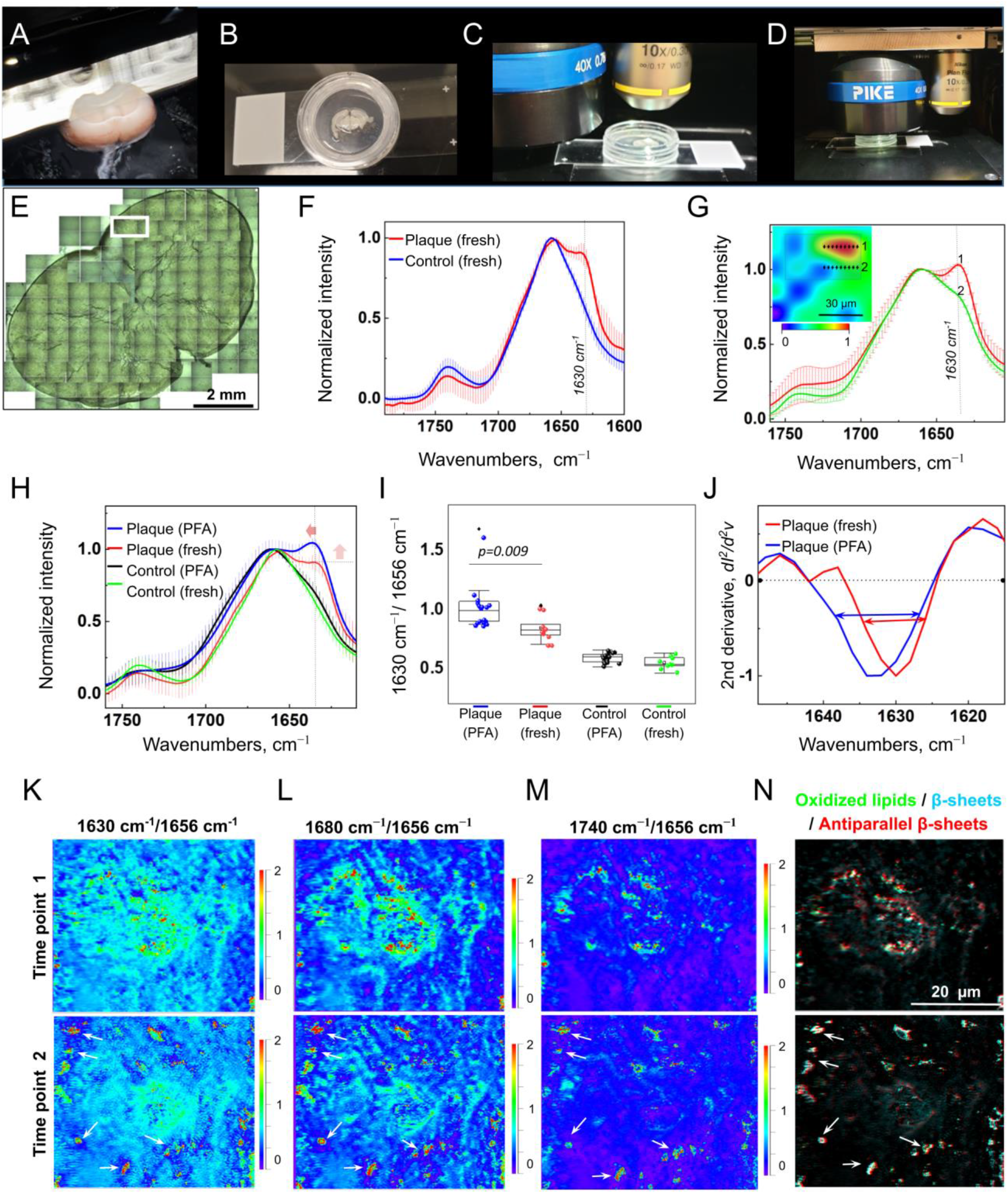
Time-resolved imaging of amyloid structures in living tissue at submicron resolution. (**A**) Vibratome-assisted slicing (275 μm thickness) of a fresh brain of AD transgenic mouse. (**B**) Incubation of tissue sections in a medium at room temperature. (**C**) Partial removal of the medium before O-PTIR analysis, allowing for tissue hydration and focusing using a 15×-air objective. (**D**) OPTIR measurements using 40 × -air objective. (**E**) Sample (same as in **Fig. 2B**) and O-PTIR measurement area (white rectangle). (**F**) Averaged and normalized O-PTIR spectra collected from several amyloid plaques and control area. (**G**) A slice of a hyperspectral map of IR absorption at 1630 cm^-1^ normalized to 1656^-1^; the insert shows spatial distribution of normalized 1630 cm^-1^ intensity. Averaged and normalized spectra correspond to the spectra line indicated by corresponding numbers. (**H**) O-PTIR analysis of fresh and tissue fixed in 4 % PFA. Arrows indicate an increase in the intensity of β-sheet (up) and changes of structural content, band position shift from 1630 cm^-1^ to 1634 cm^-1^. (**I**) β-sheet signal intensity in living and fixed tissue. Tukey’s post-hoc test; N=8–10 spots. Data are the mean ± s.d. (**J**) Second derivatives of IR absorbance spectra. Arrows indicate changes in bandwidth. (**K-N**) Single-frequency IR maps of the same spot in living tissue at time 0 and time 10 min. (**K**) Change of β-sheet structure distribution over time, calculated as a ratio of 1630 cm^-1^ (the main β-sheet band) to 1656 cm^-1^ signal (Amide I maximum). Lower panel, white arrows indicate newly formed β-sheet structures. (**L**) Change of distribution of fibrillar β-sheet structures over time, calculated as a ratio of 1680 cm^-1^ (antiparallel β-sheets) to 1656 cm^-1^ intensity. (**M**) Change in lipid oxidation over time, calculated as a ratio of 1740 cm^-1^ (R-CO-OR groups) to 1656 cm^-1^ intensity. Lower panel, white arrows indicate new spots of lipid oxidation. (N) Overlay of (K, L and J) single-frequency IR maps, white arrows indicate colocalization of newly formed antiparallel β-sheets and oxidized lipids. The scale bar is the same for (K–N). Error bars in **E** and **F,** s.d.

After confirming the feasibility of using O-PTIR for amyloid plaque analysis in living brain tissue, we tested its utility for time-resolved observation of the same amyloid plaque, which is not possible using the conventional IR techniques requiring tissue fixation and dehyrdation. For serial measurements on the same section, tissue was deposited in our custom-made sample support which allowed the objective to reach the tissue while maintaining a minimal amount of buffer around the brain slice (**Fig. 3 A-D**). We acquired and compared multiple single-frequency IR maps from the same area to record the distribution of specific chemical features in tissue over time. To visualize β-sheet structures, we acquired IR maps at the frequency 1630 cm^-1^ and a band centered at 1680 cm^-1^; a double-band feature for amyloid proteins is consistent with the accepted IR signature for antiparallel β-sheets^24,35,36^. Since oxidized lipids have been previously associated with amyloid structures,^14,22,37^ we also acquired a map at the frequency 1740 cm^-1^, corresponding to oxidized lipids. Thus, we obtained and compared 4 different spectral maps showing the distribution of chemical features at two distinct time points (2.5 min per map; 10 minutes between each spectral map collected at each time point). Strikingly, we observed the formation of new β-sheet structures over time (**Fig. 3K**) with a high content of antiparallel β-sheet structures^32,38^ (**Fig. 3L**). The appearance of new β-sheet structures were accompanied by an increase in the presence of oxidized lipids that accompany amyloid aggregation in AD tissue^22^ (**Fig. 3M**).

To further explore the potential of O-PTIR as a nondestructive technique for *in vivo* imaging, we analyzed embryos of vertebrate *Pleurodeles waltl* (**Fig. 4A**). We chose this stage as embryonic development provides a highly sensitive time window and any tissue damage would likely manifest in altered development. First, we acquired O-PTIR spectra from different regions of the developing embryo (**Fig. 4A, red insets**) and identified the main signal peaks to be at 1748, 1656, 1640, and 1470 cm^-^1 (**Fig. 4B**)^39^. During the measurements, we microscopically observed droplets on the larva skin reminiscent of oil droplets. In order to verify their chemical content, we took advantage of the high spatial resolution of O-PTIR and acquired pinpoint spectra from these droplets (**Fig. 4C and fig. S3**). Chemical analysis revealed a strong peak centered at 1748 cm^-1^, indicating a high amount of esters (-C=O,-C=OOH), characteristic of lipids^40^. Interestingly we also observed a significant peak shift from 1740 cm^-1^ to 1748 cm^-1^, potentially indicating enrichment in -C=OOH or alternatively the presence of a specific interaction that can shift the ester band 10 cm^-1^ up. (**Fig. 4B, C**). Further complementary analytical techniques and experiments are needed to provide more comprehensive information as to the content of these droplets as well as their origin. Next, we tuned the wavelength to 1748, 1656, 1640, and 1470 cm^-1^ and constructed chemical maps of the dorsal side of the embryo (**Fig. 4D, E**). Both lipid droplets and proteins were both visible on single-frequency IR maps at 1750 cm^-1^ and 1656 cm^-1^, respectively (**Fig. 4E**). Importantly, all embryos survived O-PTIR (n=7) and developed similarly to controls not exposed to O-PTIR (n=?), (**Fig. 4F**), demonstrating that the IR and light intensities used with O-PTIR do not cause tissue or organism level damage which prevents normal embryonic development and maturation. Overall, the label-free nature of O-PTIR allows for *in vivo* spectroscopic investigation of biological molecules that would otherwise require sacrificing the animal and chemical tissue processing.

**Fig. 4.**
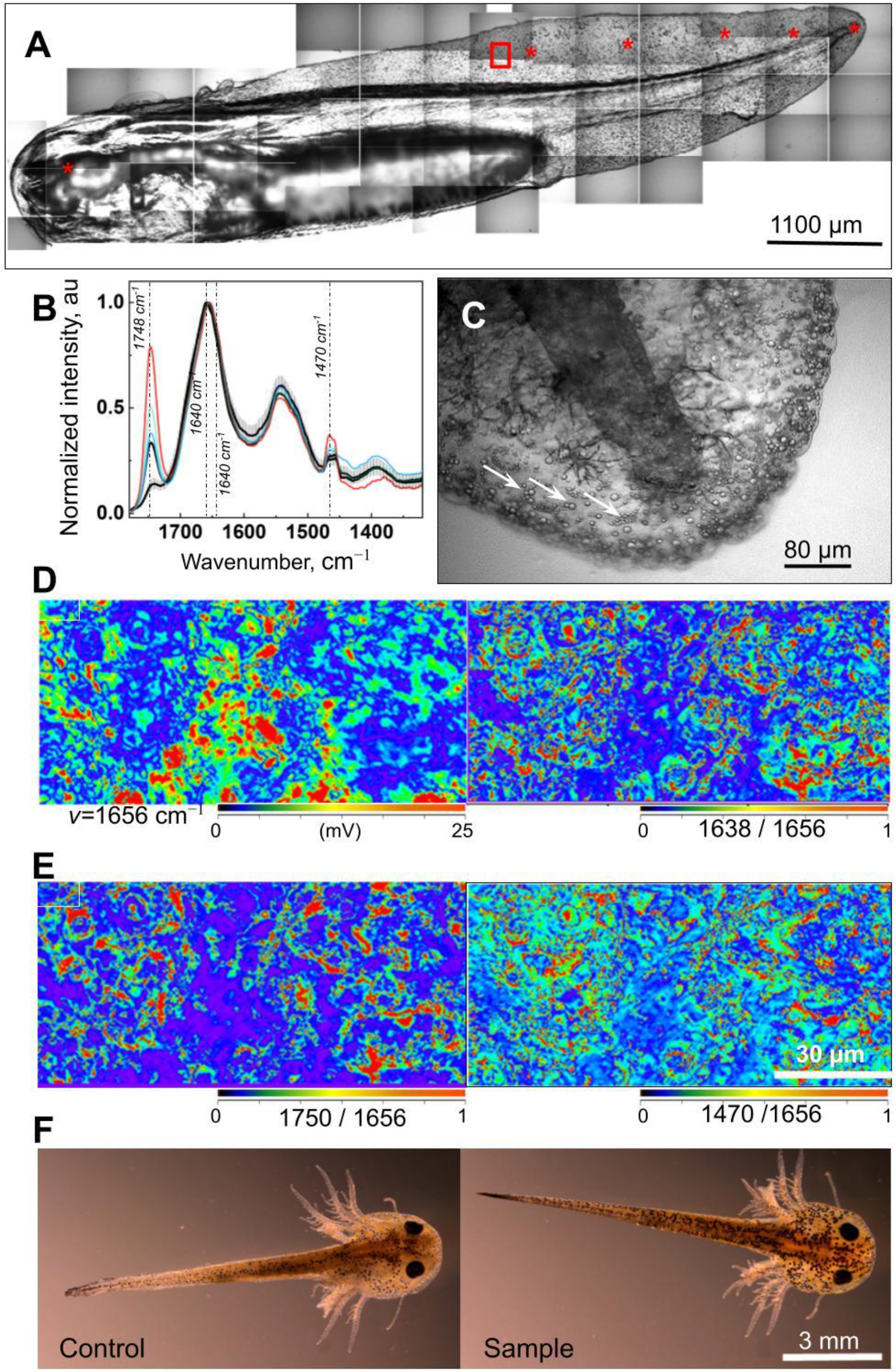
*In vivo* O-PTIR imaging of lipids and proteins in *P. waltl*. (**A**) Bright field mosaic of the *P. waltl* embryo. Asterisks indicate the O-PTIR spectrum acquisition points. Red rectangle indicates the O-PTIR map position shown in (**D**) and (**E**). (**B**) Normalized O-PTIR spectra acquired at different locations are shown in (**A**). Black traces, spectra acquired from spots outside lipid droplets; error bars, s.d. for black-trace spectra; N=7 animals. Colored traces, spectra acquired from the lipid droplets. Dashed lines show the wavenumber for single frequency IR maps in (**D**) and (**E**). (**C**) Bright field image of the embryonic tail. Arrows indicate lipid droplets on the skin. Spectra for the droplet and area outside the droplet are shown in **Fig. S2** (**D**) Single frequency IR maps acquired at the same spot at two energy levels corresponding to proteins (1656 cm ^-1^ and 1640 cm^-1^). (**E**) Single-frequency IR maps acquired at the same spot at two energy levels corresponding to R-CO-OR groups (1750 cm^-1^ and 1470 cm^-1^), showing the distribution of lipids. (**F**) Brightfield images of *P. waltl* embryos 12 d after O-PTIR measurements. Compared to the control animals not analyzed by O-PTIR, no change in animal development was apparent.

## Discussion

We demonstrate that O-PTIR can be used as a novel IR imaging platform for acquisition of time-resolved imaging of lipids and proteins in fresh biopsies of diverse tissues as well as in in living brain tissue and whole organisms with sub-micron spatial resolution^13^. Importantly, O-PTIR resolution is 10 times better than that of μIR, i.e., approximately 400 nm for O-PTIR vs. 4.6 μm for conventional μIR, at 1630 cm^-1^ corresponding to β-sheet structures and using the Rayleigh criterion of 0.61λ/NA (where λ is a wavelength, NA is the numerical aperture of the objective, here equal 0.8). This opens up the possibility to visualize biomolecular structures at submicron resolution in living tissue and for example, to monitor the onset, structural evolution and spread of amyloid β-sheet aggregates in living brain tissue. To the best of our knowledge, this is the first indication in real-time, living tissue that newly formed antiparallel β-sheet structures are accompanied by lipid changes (**Fig. 3M**). Interestingly, we observed that some antiparallel β-sheets present in the initial time point were not visible in the second time point and that these regions coincide with a decrease in lipid oxidation overtime. Although additional studies are needed to understand this phenomena, including the use of antioxidants or free radical scavengers in the culture media, we hypothesize that oxygen exposure directly or indirectly triggered the formation of new amyloid β-sheet aggregates (**Fig. S4**). Although more studies are needed, our preliminary O-PTIR experiments already revealed alteration of amyloid β-sheet structures in living tissue. We observed the formation of new antiparallel β-sheets which was accompanied by changes in the intensity of 1740 cm^-1^ which corresponds to stretching vibrations of ester bonds between phospholipids,^40^ known to be important for synaptic neurotransmission and plasticity^41^. As changes of 1680 cm^-1^, 1630 cm^-1^ along with 1740 cm^-1^ intensity indicate formation of β-sheet structures and oxidized lipids, and as a combination can be used a s biomarkers to measure *in situ* neurotoxicity, thus O-PTIR can become a valuable tool for amyloid researchers to elucidate amyloid structure-neurotoxicity relationship, which is currently debated.

As O-PTIR allows for collection of IR spectra in fresh tissue samples and living organisms, it is a promising tool for use as an *in vivo* multi-omics tool for obtaining comprehensive information on various sub-cellular components (proteins, lipids, metabolites, DNA, RNA, etc.). Furthermore, as biologically molecules are highly conserved across evolution, O-PTIR opens up new opportunities for biological tissues and organisms which currently have limited research tools to (e.g. model organisms such as *P. waltl* or research species with limited species-specific antibodies). Lipid droplets on *P. waltl* embryonic skin have not been reported to date and thus their function is presently unknown. They may act as nutrient stores during embryonic development in addition to the yolk sac or alternatively serve to secrete metabolites in a sequestered format to prevent subsequent uptake by the developing organism. Further studies are needed to understand the biological importance of this finding as well as further studies to better understand their specific composition. Thus, we show that O-PTIR is well-tolerated by developing *P. waltl* embryos and is therefore a promising new techniques to study live animals in a label-free manner, with no need for extensive sample processing.

In the future, an exciting potential area of application for O-PTIR imaging will be for rapid molecule-based diagnosis of clinical diseases. For instance, on-site O-PTIR analysis could be used to rapidly screen biopsies during cancer surgery, such as Mohs surgery for skin cancer^42^, substantially cutting down on the analysis as well as surgical time. Other potential clinical applications will rely on further instrument development. There is a huge clinical need for techniques which can be used to evaluate intact organs without taking a biopsy (e.g. evaluation of donor organs prior to transplantation). The application of O-PTIR to whole, intact organs or larger organisms would require further technical developments such as an adjustable sample stage or flexible or portable optics for the analysis of samples larger than a few centimeters thick.

Notwithstanding our promising findings, O-PTIR can be time-consuming as a standalone technique which may make it suboptimal for assessment of some living samples. Further instrument and accessory developments are thus needed to overcome this limitation. As one example, instruments combining O-PTIR and epifluorescence modules could be used to decrease the imaging acquisition time as specific antibodies for immunolabeling or transgenic animals containing fluorescent proteins could be used to rapidly locate immunolabeled areas. In addition to a reduction in acquisition time, such a development could also add specificity for spectroscopic measurements.

In summary, we demonstrate that OPTIR using modified imaging acquisition parameters allows it to be used as a novel IR imaging platform to provide sub-micron spatial resolution, enabling imaging of biological molecules such as lipids and proteins in hydrated, living tissues and organisms, as well as insights into their structural conformations. As a proof-of-concept, we conducted multiple experiments that demonstrated the utility of O-PTIR as a novel multi-omic approach to record spectra directly from multiple tissues. Overall, O-PTIR imaging promises broad applications that go beyond the reach of current IR microscopy approaches, with implications for such diverse fields as basic biomedical research, proteinopathies such as Alzheimer’s disease research, or medical diagnosis of biochemical changes in tissues.

## Methods

### Preparation of mouse brain slices

All mouse experiments were compliant with the requirements of the Ethical Committee of Lund University (M11908-19). APP/PS1 mice (hAPPswe, PSEN1dE9)85Dbo/Mmjax were obtained from Jackson Labs, USA, and all mice were screened for the presence of the human APP695 transgene by PCR.

Briefly, the animals were anaesthetized intraperitoneally with sodium pentobarbital (40–50 mg/kg) and transcardially perfused with ~20 mL of oxygenated (carbogen, 5 % CO2, 95 % O_2_) N-methyl-d-glucamine (NMDG)-HEPES artificial cerebrospinal fluid (CSF) containing 92 mM NMDG, 2.5 mM KCl, 1.25 mM NaH_2_PO_4_, 30 mM NaHCO_3_, 20 mM HEPES, 25 mM glucose, 2 mM thiourea, 5 mM Na-ascorbate, 3 mM Na-pyruvate, 0.5 mM CaCl_2_·2H_2_O, and 10 mM MgSO_4_·7H_2_O. pH was tittrated to 7.3–7.4 with 7 mL ± 0.2 mL of 37 % hydrochloric acid, with osmolarity ranging from 300 to 305 mOsm/kg. The brain was quickly removed from the skull and placed in an ice-cold NMDG-HEPES buffer. Acute brain slices containing perirhinal cortex and hippocampus were prepared at 275 μm thickness using a vibratome (Leica VT1200 S, Wetzlar, Germany). Slices were incubated for 25 min at 35° C and then transferred at room temperature into a custom made holding chamber.

### Electrophysiology

Ex vivo whole-cell patch-clamp recordings were carried out on mouse brain slices. Neuronal cells were visualized using a fixed-stage Olympus Microscope (BX51WI) coupled with an IR-CCD camera and a 40X water immersion objective and recorded with continuous perfusion of artificial cerebrospinal fluid (aCSF, 1 ml/min) containing 124 mM NaCl, 2.5 mM KCl, 1.25 mM NaH_2_PO_4_, 24 mM NaHCO_3_, 12.5 mM glucose, 5 mM HEPES, 2 mM CaCl_2_·2H_2_O, and 2 mM MgSO_4_·7H_2_O, pH 7.4; 305 mOsm/kg. Borosilicate glass electrodes (4–6 MΩ resistance) were filled with intracellular solution containing: 130 mM (K) Gluconate, 10 mM KCl, 0.2 mM EGTA, 10 mM HEPES, 4 mM (Mg)ATP, 0.5 mM (Na)GTP, 10 mM (Na)Phosphocreatine (pH 7.25, 296 mOsm). Recordings were obtained using a Multiclamp 700B amplifier and pClamp 10.4 data acquisition software (Axon Instruments, Molecular Devices, USA). Original traces were obtained offline using Clampfit 10.6 (Axon Instruments, Molecular Devices, USA)

### Pleurodeles waltl imaging

Embryonic wild-type *Pleurodeles waltl* (stage 34-36, 12 days post fertilization (dpf)) used in this study were from the colony at Lund University. Swedish regulations (Jordbruksverkets föreskrift L150, §5) state that working with vertebrate embryos before their ability to feed independently does not require Institutional Animal Care and Use Committee oversight and the stage selected is before this independent feeding stage. Embryos were reared in conditioned tap water at 20-22°C (55g Tetra marine salt, 15g of Ektozon salt, and 2.5ml of Prime^™^ water de-chlorinator per 100 litres of water). For imaging, embryos were removed from their protective jelly using fine-tip forceps at 10 dpf. At 12 dpf, corresponding to stage 34-36^43^, embryos were narcotized in 0.025 % of tricaine in conditioned tap water (pH 7.0). Once completely narcotized, embryos were then moved via a transfer pipette to a microscope slide for imaging. After imaging, embryos were then transferred back to the container with conditioned tap water (pH 7.0), all embryos were allowed to develop into larvae to assess for morphological damage.

### Spectroscopy

O-PTIR imaging was performed at Lund (Integrated vibration spectroscopy - microcosm laboratory for molecular-scale biogeochemical research, Lund University). The IR source was a pulsed, tunable four-stage QCL device, scanning from 1800 to 800 cm^-1^ at 100 kHz repetition rate. The probe was a CW 532 nm visible variable power laser. The photothermal effect was detected through the modulation of the green laser intensity induced by the pulsed IR laser. Further details about the fundamentals of the technique and the instrument itself can be found in references^44,45^. Spectra were averaged for 5–8 scans. The collection parameters were: spectral range 1790–1000 cm^-1^, reflection mode at 2 cm^-1^ spectral resolution. To avoid photodamage, that has been described for O-PTIR^46^, IR power was 22 %, green light probe power was 6 %, pulse rate was set to 100 kHz, Mirage DC was 1.005 V, and 10x standard detector gain. Background spectra were collected on an in-built reference sample. O-PTIR spectra were normalized to the max at 1656 cm^-1^; second-order derivation of the spectra was used to increase the number of discriminative features; the Savitsky-Golay algorithm with a 5-point filter with 3rd polynomial order was employed in this process. O-PTIR absorption images of living brain tissue were obtained with a linear scan speed of 100 μm/sec with total acquisition times in the range of 2-3 minutes for 45 μm x 37 μm with the step of 250 nm. O-PTIR absorption images of embryos were obtained with a linear scan speed of 100 μm/sec with total acquisition times in the range of 5 minutes for 100 μm x 50 μm with the step of 150 nm. The relative fraction of β-sheet structures was visualized by calculating the map intensity ratio between 1630 cm^-1^, a peak corresponding to β-sheet structures^1,24,32^and 1656 cm^-1^, maximum of Amide I.^1^ The increase of intensity in the resultant ratio map was considered a sign of the higher concentration of amyloid fibrils. The intensity at 1740 cm^-1^ was considered a sign of the higher concentration of esters^40^.

### Tissue fixation and processing

After brain extraction, the animal’s chest and abdominal cavities were opened and all organs were removed and placed on petri dish on ice. Approx. 2 x 1.5 mm tissue biopsies were obtained using a puncher (WellTech, Denmark) and placed in 96 well plate containing cell culture media. After spectroscopy and viability assay (described below), tissue was fixed overnight at 4° C in 10 % neutral buffer formalin solution (Sigma-Aldrich, Sweden) and processed as previously described^47^. Specifically, formalin-fixed tissue was passed through graded ethanol (Solveco, Sweden) and isopropanol (Univar Solutions, Sweden) solutions prior to paraffin embedding (Histolab, Sweden) in an automated spin tissue processor (Myr, Spain) using processing/embedding cassettes (Histolab, Sweden)^47^. Paraffin-embedded tissue biopsies were cut into 4 μm tissue sections using a microtome (Litz,Germany) and mounted on super frost adhesive microscope slides (Epredia, USA). Before chemical de-paraffinization, slides were placed overnight in a 65° C oven in a horizontal position, followed by de-paraffinization using Histo-Clear (Ted Pella, USA). After de-paraffinization, tissue slides were stained with Hematoxylin and Eosin (Merck Millipore, Germany), dehydrated, mounted with Pertex Mounting media (Histolab, Sweden) and imaged using VS120-S6-096 virtual microscopy slide scanning system (Olympus, Tokyo, Japan).

### Tissue Imaging

A fluorescent image of rehydrated brain tissue with a thickness of 275 μm stained with Amy tracker and placed on glass microscopic slides was acquired using the manual mode in a Cytation 5 multimode reader (Agilent Technologies, USA). Montage images were collected at 4x and 20x magnification. Texas red and GFP imaging filter cubes were used with the following acquisition settings: GFP LED intensity: 4, integration time: 5 msec, camera gain: 22. Texas red: LED intensity: 4, integration time: 16 msec, camera gain: 24, both with brightness 50 units and contrast 33 units.

### Confocal microscopy

After O-PTIR imaging, the presence of amyloid plaques in fresh APP/PS1 brain tissue was confirmed by staining for Amytracker^R^ 520 (EbbaBiotech, Solna, Sweden). A laser scanner Leica TCS SP8 confocal microscope (Leica Microsystems) with Leica Application Suite X software (version 3.4.7.23225), was used to obtain high-resolution images of amyloid plaques. 5x air and 40x oil objectives in combination with a 568 laser and a z-step size of 1.0 μm were used to acquire the fluorescence images.

### Water-soluble tetrazolium-1viability assay (WST-1)

Extracellular reduction of water-soluble tetrazolium salt reagent (WST-1) was used to assess tissue viability^48^. Brain slices or biopsy pieces were incubated with 100 μL DMEM/F12 (Gibco, Sweden) cell culture media supplemented with 0.1 % FBS (Gibco, Sweden), 1 % penicillin/streptomycin, (Gibco, USA), 1 % AmpB (Amphotericin B, Gibco, UK) and 10 μL WST-1 (Roche, Sigma-Aldrich, USA) for 1 hour at 37° C in a humidified incubator with 5% CO_2_. Supernatant optical density was measured at 440 and 650 nm on an Epoch plate reader (BopTeck, USA). 440nm represents the specific conversion of WST-1 by metabolically active cells and 650 nm wavelength was used as a reference wavelength for non-specific absorbance. Complete mediums used for each tissue served as negative controls.

## Acknowledgments

The authors are indebted to Prof. Gunnar K. Gouras for the APP/PS1 animals. The authors acknowledge Elevate Scientific for the editorial help.

## Author contributions

Conceptualization: DEW, NDL, OK

Methodology: NG, VS, IAS, ECP, SK, OK, CHU, DEW, JD

Visualization: NG, VS, IAS, SK

Funding acquisition: DRO, DEW, NDL, OK

Project administration: NDL, OK

Supervision: DEW, NDL, OK

Writing – original draft: NG, ES, ECP, SK

Writing – review & editing: DEW, NDL, OK

## Data and materials availability

All data and materials used in the analysis are available in some form to any researcher for purposes of reproducing or extending the analysis.

## Supplementary Materials

Materials and Methods

Supplementary Figs. S1-4

References (*46*)

## References and Notes

1. Barth, A. Infrared spectroscopy of proteins. Biochimica et Biophysica Acta (BBA) - Bioenergetics 1767, 1073–1101 (2007).

2. Naumann, D. & Diem, M. Modern biophotonic trends in microbiological and medical diagnostics. J. Biophoton. 3, 492–492 (2010).

3. Klementieva, O. et al. Pre-plaque conformational changes in Alzheimer’s disease-linked Aβ and APP. Nature Communications 8, 14726 (2017).

4. Old, O. J. et al. Vibrational spectroscopy for cancer diagnostics. Analytical Methods 6, 3901 (2014).

5. Albert, J.-D. et al. A novel method for a fast diagnosis of septic arthritis using mid infrared and deported spectroscopy. Joint Bone Spine 83, 318–323 (2016).

6. Roy, S., Perez-Guaita, D., Bowden, S., Heraud, P. & Wood, B. R. Spectroscopy goes viral: Diagnosis of hepatitis B and C virus infection from human sera using ATR-FTIR spectroscopy. Clinical Spectroscopy 1, 100001 (2019).

7. Agbaria, A. H. et al. Rapid diagnosis of infection etiology in febrile pediatric oncology patients using infrared spectroscopy of leukocytes. Journal of Biophotonics 13, (2020).

8. Zhang, D. et al. Depth-resolved mid-infrared photothermal imaging of living cells and organisms with submicrometer spatial resolution. Sci. Adv. 2, e1600521 (2016).

9. Kansiz, M. et al. Review of life science applications using submicron O-PTIR and simultaneous Raman microscopy: a new paradigm in vibrational spectroscopy. in Advanced Chemical Microscopy for Life Science and Translational Medicine 2021 (eds. Simpson, G. J., Cheng, J.-X. & Min, W.) 10 (SPIE, 2021). doi:10.1117/12.2578662.

10. Lima, C., Muhamadali, H., Xu, Y., Kansiz, M. & Goodacre, R. Imaging Isotopically Labeled Bacteria at the Single-Cell Level Using High-Resolution Optical Infrared Photothermal Spectroscopy. Anal. Chem. 93, 3082–3088 (2021).

11. Su, Y. et al. Steam disinfection releases micro(nano)plastics from silicone-rubber baby teats as examined by optical photothermal infrared microspectroscopy. Nat. Nanotechnol. 17, 76–85 (2022).

12. Rahmati, M. et al. Intrinsically disordered peptides enhance regenerative capacities of bone composite xenografts. Materials Today 52, 63–79 (2022).

13. Klementieva, O. et al. Super-Resolution Infrared Imaging of Polymorphic Amyloid Aggregates Directly in Neurons. Adv. Sci. 1903004 (2020) doi:10.1002/advs.201903004.

14. Gustavsson, N. et al. Correlative optical photothermal infrared and X-ray fluorescence for chemical imaging of trace elements and relevant molecular structures directly in neurons. Light Sci Appl 10, 151 (2021).

15. Paulus, A. et al. Correlative imaging to resolve molecular structures in individual cells: Substrate validation study for super-resolution infrared microspectroscopy. Nanomedicine: Nanotechnology, Biology and Medicine 43, 102563 (2022).

16. Kansiz, M. et al. Optical Photothermal Infrared Microspectroscopy Discriminates for the First Time Different Types of Lung Cells on Histopathology Glass Slides. Anal. Chem. 93, 11081–11088 (2021).

17. Bakir, G. et al. Orientation Matters: Polarization Dependent IR Spectroscopy of Collagen from Intact Tendon Down to the Single Fibril Level. Molecules 25, 4295 (2020).

18. Spadea, A., Denbigh, J., Lawrence, M. J., Kansiz, M. & Gardner, P. Analysis of Fixed and Live Single Cells Using Optical Photothermal Infrared with Concomitant Raman Spectroscopy. Anal. Chem. acs.analchem.0c04846 (2021) doi:10.1021/acs.analchem.0c04846.

19. Malek, K., Wood, B. R. & Bambery, K. R. FTIR Imaging of Tissues: Techniques and Methods of Analysis. in Optical Spectroscopy and Computational Methods in Biology and Medicine (ed. Baranska, M.) vol. 14 419–473 (Springer Netherlands, 2014).

20. Le Naour, F. et al. Chemical Imaging on Liver Steatosis Using Synchrotron Infrared and ToF-SIMS Microspectroscopies. PLoS ONE 4, e7408 (2009).

21. Szafraniec, E. et al. Vibrational spectroscopy-based quantification of liver steatosis. Biochimica et Biophysica Acta (BBA) - Molecular Basis of Disease 1865, 165526 (2019).

22. Benseny-Cases, N., Klementieva, O., Cotte, M., Ferrer, I. & Cladera, J. Microspectroscopy (μFTIR) reveals co-localization of lipid oxidation and amyloid plaques in human Alzheimer disease brains. Anal. Chem. 86, 12047–12054 (2014).

23. Orphanou, C.-M. The detection and discrimination of human body fluids using ATR FT-IR spectroscopy. Forensic Science International 252, e10–e16 (2015).

24. Cerf, E. et al. Antiparallel β-sheet: a signature structure of the oligomeric amyloid β-peptide. Biochemical Journal 421, 415–423 (2009).

25. Aso, E. et al. Amyloid Generation and Dysfunctional Immunoproteasome Activation with Disease Progression in Animal Model of Familial Alzheimer’s Disease: Amyloid Generation and UPS in FAD Mice. Brain Pathology 22, 636–653 (2012).

26. Nilsson, K. P. R., Herland, A., Hammarström, P. & Inganäs, O. Conjugated Polyelectrolytes: Conformation-Sensitive Optical Probes for Detection of Amyloid Fibril Formation. Biochemistry 44, 3718–3724 (2005).

27. Ting, J. T. et al. Preparation of Acute Brain Slices Using an Optimized N-Methyl-D-glucamine Protective Recovery Method. JoVE 53825 (2018) doi:10.3791/53825.

28. Elkins, M. R. et al. Structural Polymorphism of Alzheimer’s β-Amyloid Fibrils as Controlled by an E22 Switch: A Solid-State NMR Study. Journal of the American Chemical Society 138, 9840–9852 (2016).

29. Kollmer, M. et al. Cryo-EM structure and polymorphism of Aβ amyloid fibrils purified from Alzheimer’s brain tissue. Nat Commun 10, 4760 (2019).

30. Fändrich, M., Meinhardt, J. & Grigorieff, N. Structural polymorphism of Alzheimer Aβ and other amyloid fibrils. Prion 3, 89–93 (2009).

31. Byler, D. M. & Susi, H. Examination of the secondary structure of proteins by deconvolved FTIR spectra. Biopolymers 25, 469–487 (1986).

32. Vosough, F. & Barth, A. Characterization of Homogeneous and Heterogeneous Amyloid-β42 Oligomer Preparations with Biochemical Methods and Infrared Spectroscopy Reveals a Correlation between Infrared Spectrum and Oligomer Size. ACS Chem. Neurosci. 12, 473–488 (2021).

33. Swanson, C. J. et al. A randomized, double-blind, phase 2b proof-of-concept clinical trial in early Alzheimer’s disease with lecanemab, an anti-Aβ protofibril antibody. Alz Res Therapy 13, 80 (2021).

34. Schneider, L. A resurrection of aducanumab for Alzheimer’s disease. The Lancet Neurology S1474442219304806 (2019) doi:10.1016/S1474-4422(19)30480-6.

35. Sarroukh, R., Goormaghtigh, E., Ruysschaert, J.-M. & Raussens, V. ATR-FTIR: A “rejuvenated” tool to investigate amyloid proteins. Biochimica et Biophysica Acta (BBA) - Biomembranes 1828, 2328–2338 (2013).

36. Li, H., Lantz, R. & Du, D. Vibrational Approach to the Dynamics and Structure of Protein Amyloids. Molecules 24, 186 (2019).

37. Arimon, M. et al. Oxidative stress and lipid peroxidation are upstream of amyloid pathology. Neurobiology of Disease 84, 109–119 (2015).

38. Zou, Y., Li, Y., Hao, W., Hu, X. & Ma, G. Parallel β-Sheet Fibril and Antiparallel β-Sheet Oligomer: New Insights into Amyloid Formation of Hen Egg White Lysozyme under Heat and Acidic Condition from FTIR Spectroscopy. J. Phys. Chem. B 117, 4003–4013 (2013).

39. Aragão, B. J. G. de & Messaddeq, Y. Peak separation by derivative spectroscopy applied to ftir analysis of hydrolized silica. Journal of the Brazilian Chemical Society 19, 1582–1594 (2008).

40. Dreissig, I., Machill, S., Salzer, R. & Krafft, C. Quantification of brain lipids by FTIR spectroscopy and partial least squares regression. Spectrochimica Acta Part A: Molecular and Biomolecular Spectroscopy 71, 2069–2075 (2009).

41. García-Morales, V. et al. Membrane-Derived Phospholipids Control Synaptic Neurotransmission and Plasticity. PLoS Biol 13, e1002153 (2015).

42. Golda, N. & Hruza, G. Mohs Micrographic Surgery. Dermatologic Clinics 41, 39–47 (2023).

43. Shi, D. L. & Boucaut, J. C. The chronological development of the urodele amphibian Pleurodeles waltl (Michah). Int J Dev Biol 39, 427–441 (1995).

44. DeVetter, B. M., Kenkel, S., Mittal, S., Bhargava, R. & Wrobel, T. P. Characterization of the structure of low-e substrates and consequences for IR transflection measurements. Vibrational Spectroscopy 91, 119–127 (2017).

45. Kansiz, M. et al. Optical Photothermal Infrared Microspectroscopy with Simultaneous Raman – A New Non-Contact Failure Analysis Technique for Identification of <10 μm Organic Contamination in the Hard Drive and other Electronics Industries. Microscopy Today 28, 26–36 (2020).

46. Sandt, C. & Borondics, F. Super-resolution infrared microspectroscopy reveals heterogeneous distribution of photosensitive lipids in human hair medulla. Talanta 254, 124152 (2023).

47. Silva, I. et al. A Semi-quantitative Scoring System for Green Histopathological Evaluation of Large Animal Models of Acute Lung Injury. BIO-PROTOCOL 12, (2022).

48. Alsafadi, H. N. et al. An ex vivo model to induce early fibrosis-like changes in human precision-cut lung slices. American Journal of Physiology-Lung Cellular and Molecular Physiology 312, L896–L902 (2017).

