## Supplemental for "Label-free high-resolution infrared spectroscopy for spatiotemporal analysis of complex living systems"

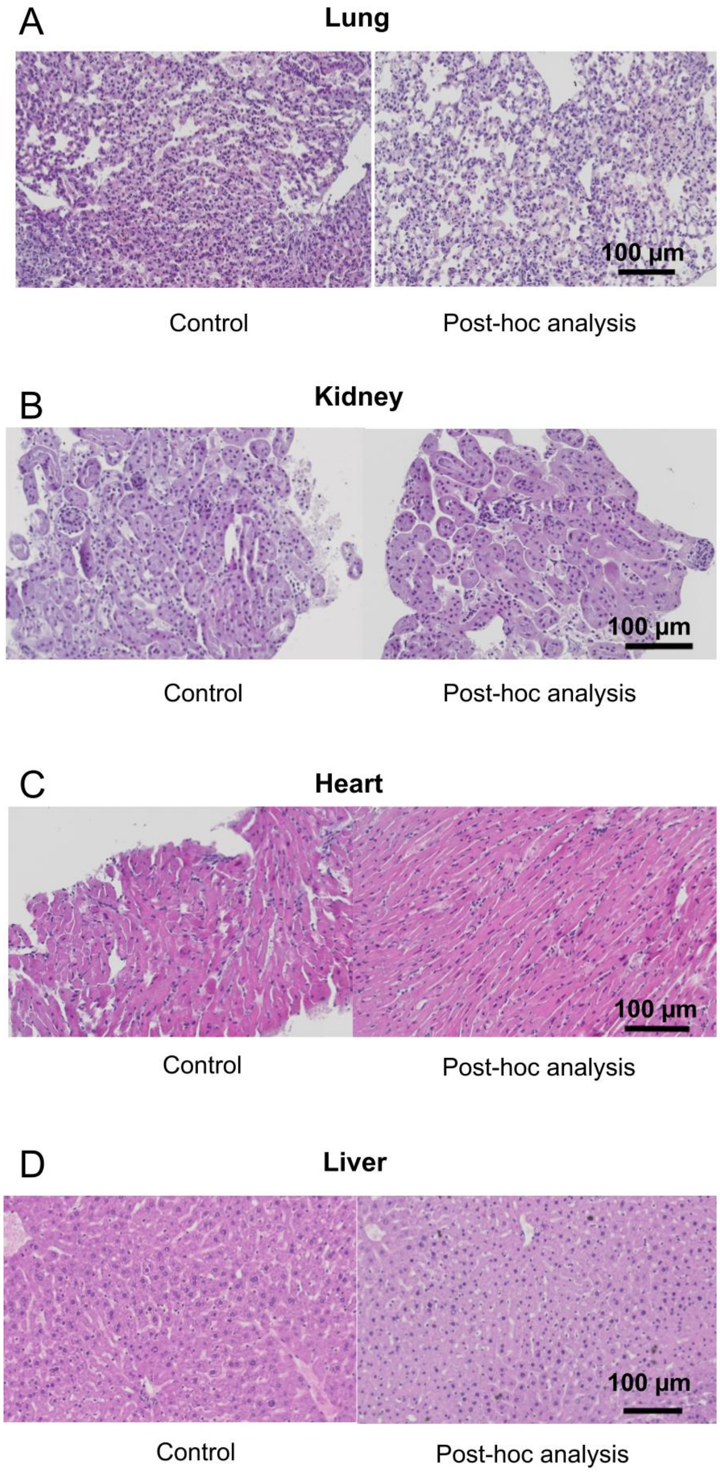


**Figure S1.** Hematoxylin-eosin stained tissue of control biopsies that were not analyzed by O-PTIR (Control) and tissue biopsies after O-PTIR (Post-hoc analysis)

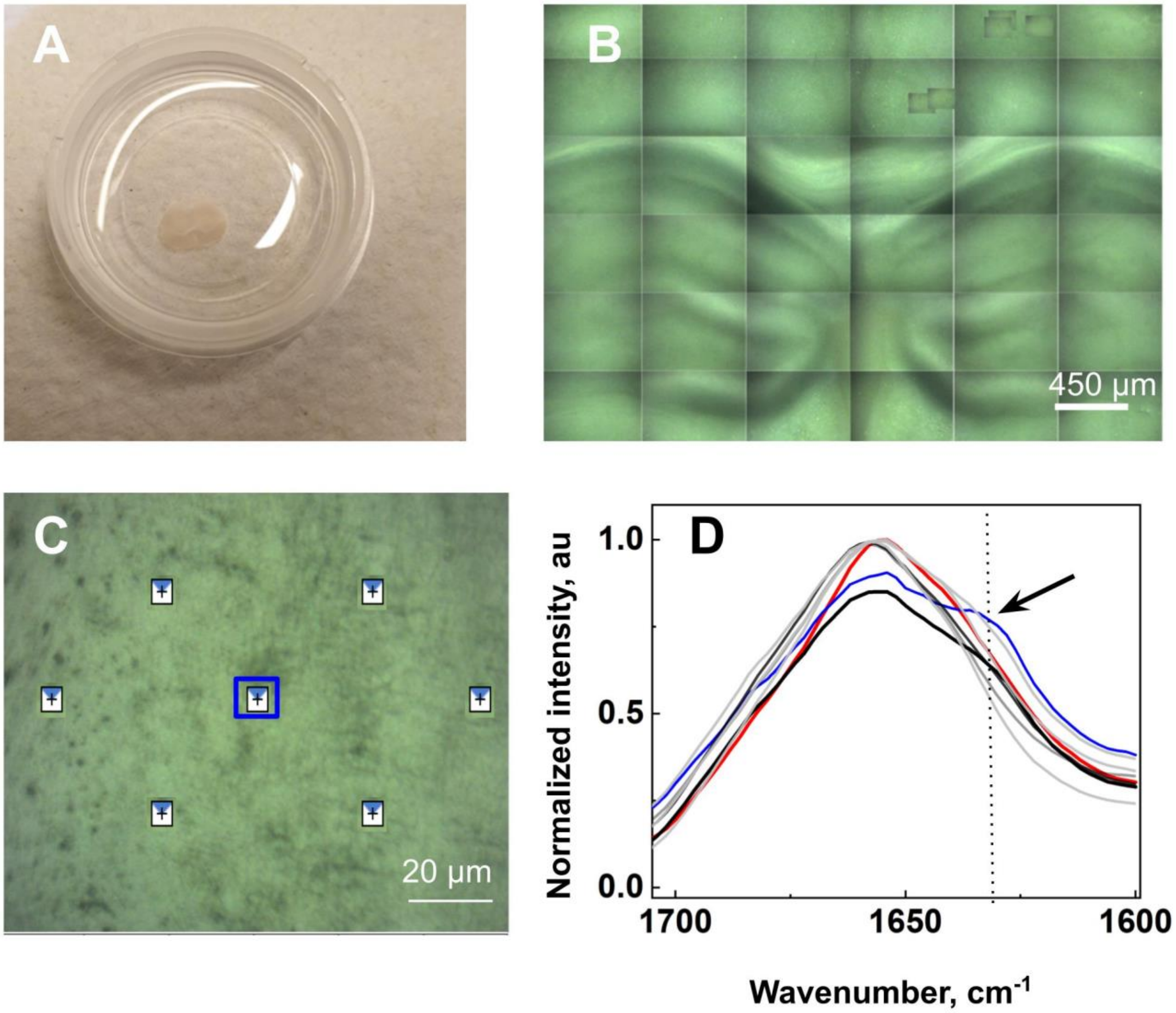


**Figure S2.** (**A**) Sample overview. (**B**) Cortex and hippocampus overview. (**C**) Example of the scanned area. (**D**) amyloid plaques were located in brain tissue using fast hyperspectral array scanning over an extended area (here is a part of the area scanned) 1 second per spectra, over 1600 $cm^{-1}$ -1700 $cm^{-1}$), as shown by an arrow, plaque was located using the associated IR absorption peak at 1630 $cm^{-1}$.

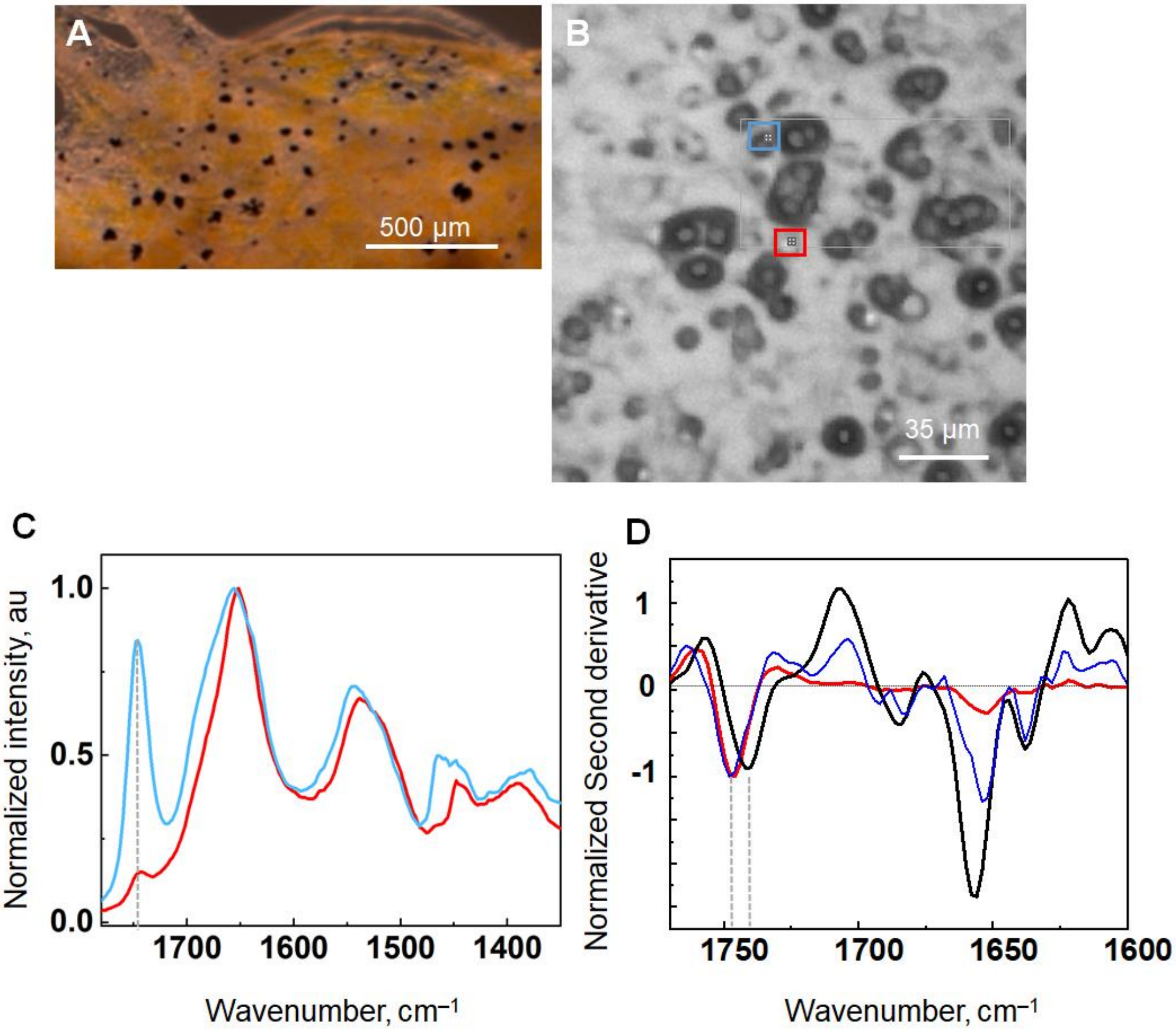


**Figure S3.** OPYIT on *Pleurodeles waltl.* (**A**) Sample overview of the *Pleurodeles waltl* embryonic skin. (**B**) Spectra locations acquired from the droplets on the *Pleurodeles waltl* embryonic skin. (**C**) Normalized O-PTIR spectra. Dashed line indicated the band located at 1750 $cm^{-1}$ corresponding to ester −C═O, −C═OOH groups. (**D**) Normalized second derivetive to define the peak positions. Black spectra brain tissue, peak centered arounf 1740 $cm^{-1}$; red spectra is dried *Pleurodeles waltl* skin; blue spectra from living *Pleurodeles waltl*. Dashed lines indicte peak position.

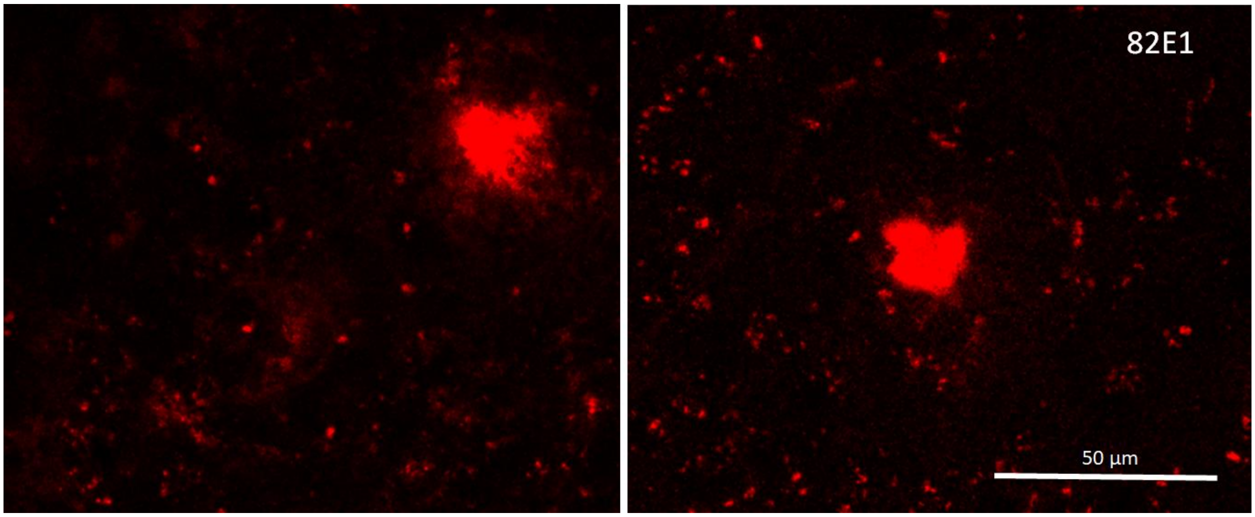


**Figure S4.** Fluorescent image of amyloid plaques in brain tissue showing amyloid proteins immunolabeled with Aβ specific antibody (82E1, red). Left panel: tissue fixed immediately after cutting. Right panel: tissue fixed after spectroscopy.
